# Comparison of commercial DNA extraction kits for whole metagenome sequencing of human oral, vaginal, and rectal microbiome samples

**DOI:** 10.1101/2023.02.01.526597

**Authors:** Michelle L. Wright, Jessica Podnar, Kayla D. Longoria, Tien C. Nguyen, Sungju Lim, Sarina Garcia, Dennis Wylie

## Abstract

**Introduction:** Advancements in DNA extraction and sequencing technologies have been fundamental in deciphering the significance of the microbiome related to human health and pathology. Whole metagenome shotgun sequencing (WMS) is gaining popularity in use compared to its predecessor (i.e., amplicon-based approaches). However, like amplicon-based approaches, WMS is subject to bias from DNA extraction methods that can compromise the integrity of sequencing and subsequent findings. The purpose of this study was to evaluate systematic differences among four commercially available DNA extraction kits frequently used for WMS analysis of the microbiome.

**Methods:** Oral, vaginal, and rectal swabs were collected in replicates of four by a healthcare provider from five participants and randomized to one of four DNA extraction kits. Two extraction blanks and three replicate mock community samples were also extracted using each extraction kit. WMS was completed with NovaSeq 6000 for all samples. Sequencing and microbial communities were analyzed using nonmetric multidimensional scaling and compositional bias analysis.

**Results:** Extraction kits differentially biased the percentage of reads attributed to microbial taxa across samples and body sites. The PowerSoil Pro kit performed best in approximating expected proportions of mock communities. While HostZERO was biased against gram-negative bacteria, the kit outperformed other kits in extracting fungal DNA. In clinical samples, HostZERO yielded a smaller fraction of reads assigned to *Homo sapiens* across sites and had a higher fraction of reads assigned to bacterial taxa compared to other kits. However, HostZERO appears to bias representation of microbial communities and demonstrated the most dispersion by site, particularly for vaginal and rectal samples.

**Conclusions:** Systematic differences exist among four frequently referenced DNA extraction kits when used for WMS analysis of the human microbiome. Consideration of such differences in study design and data interpretation is imperative to safeguard the integrity of microbiome research and reproducibility of results.

## Introduction

The diverse composition of the microbiome throughout the body plays an important role in human health. The advancement of sequencing technologies has made the assessment of microbiome composition possible through microbial DNA extraction and sequencing. Early studies of the human microbiome largely described bacterial communities using the 16S amplicon-based sequencing approach [1]. However, this amplicon-based approach does not allow for the discovery of fungi or viruses within a sample or identification of microbial genes for functional relevance such as antimicrobial resistance genes. Furthermore, results from amplicon based approaches can be biased by numerous factors including: varied microbial genetic diversity in common 16S regions, multiple 16S genome copy numbers in bacterial genomes, and choice of DNA extraction method [2]. To counter limitations of amplicon-based sequencing, there is increasing use of whole metagenome shotgun sequencing (WMS) to analyze human microbiome communities related to health outcomes.

WMS may also be impacted by the same limitations and biases as 16S amplicon-based approaches. For example, DNA extraction is crucial to the success of microbiome sequencing and is known to introduce bias in 16S studies [3–6]. The wide variations of cell membrane structures and composition can pose significant challenges to efficiency and bias-free extraction of genomic DNA. Moreover, WMS studies investigating the human microbiome can produce a disproportionately large amount of sequencing reads attributed to the human host, instead of microbes, which limits the sensitivity of the WMS approach to detect low abundance microbial species [7]. Therefore, the selection of a reliable method for DNA extraction is of paramount importance to ensure a high DNA yield and a representative characterization of microbial communities in any given body site.

The purpose of this study was to evaluate if systematic differences occur when different commercially available DNA extraction kits are used for WMS analysis of the human microbiome.

## Methods

### Ethics approval

This study was approved and monitored by the University of Texas Institutional Review Board (protocol #2019-08-0027) and the Institutional Biosafety Committee (protocol #IBC-2019-00216). Informed consent was obtained from all participants before the collection of oral, vaginal, and rectal swab samples used for this study.

### Sample collection

Participants were recruited from the Women’s Health Integrated Practice Unit (IPU) at UT Health Austin. Individuals presenting to the IPU for pelvic floor physical therapy were approached and asked if they would be interested in participating in the study. If individuals expressed interest, written informed consent was obtained. No additional information was collected about the participants or their health history. Each of five participants provided a total of 12 samples at the sample collection visit. The healthcare provider collected oral, vaginal, and rectal samples in replicates of four using sterile flocked swabs (COPAN diagnostics, Murrieta CA), in the order listed. The samples were pre-labeled by site and order of collection (e.g., oral 1, oral 2, oral 3, oral 4, etc.). The provider collecting the samples changed gloves between each site (i.e., oral, vaginal, rectal). Oral swabs were collected by first swabbing the tongue, then the roof of the mouth, and lastly around the gum line (top and bottom) twice. Vaginal samples were obtained by inserting the swab into the vagina 3-4 inches avoiding contact with the outside skin and then rotating the swab 3-4 times. Rectal samples were obtained by inserting the swab 1 inch into the rectum and then rotating the swab 3-4 times. The samples were immediately transported to the laboratory and stored at -80□ until DNA extraction was completed.

### DNA extraction

Patient samples were randomized across extraction kits using the website randomizer.org to ensure the order of swab collection did not bias the results (**Table 1**). For example, for Participant A all swabs that were collected first and labeled number one for each site (oral, vaginal, and rectal) were extracted with the HostZERO kit, the second swab collected for each site was extracted with the PowerSoil Pro kit, and so on. We used four commercially available extraction kits frequently referenced in the metagenomic literature: Qiagen DNeasy PowerSoil Pro kit (catalog number 47014), HostZERO Microbial DNA kit (catalog number D4310), PureLink Microbiome kit (catalog number A29790), and Qiagen DNeasy Blood and Tissue kit (catalog number 69504).

**Table 1.** Randomization of samples across extraction kit type.

| Extraction Kit | Participant |  |  |  |  |
| --- | --- | --- | --- | --- | --- |
|  | A | B | C | D | E |
| PowerSoil Pro | 2 | 1 | 3 | 1 | 4 |
| HostZERO | 1 | 4 | 4 | 3 | 2 |
| PureLink | 4 | 3 | 2 | 2 | 1 |
| DNeasy | 3 | 2 | 1 | 4 | 3 |

All samples using the same protocol and kit were extracted by MLW on the same day. Extraction protocols were followed per manufacturer instructions, and each protocol was completed on separate days to minimize any potential cross-contamination. All samples (i.e., oral, vaginal, rectal) that corresponded with the order number were used for the same kit. Two extraction blanks of sterile DNAse, RNAse free water (Fisher BioReagentsTM, Pittsburgh, PA) and three replicate mock community samples (ZymoBIOMICSTM mock microbial community standards catalog number D3600, Irvine, CA) were also extracted using each extraction kit at the time DNA was extracted from clinical samples. The mock communities are composed of eight bacteria [3 Gram-negative (*Pseudomonas aeruginosa, Escherichia coli, and Salmonella enterica)* and 5 Gram-positive (*Lactobacillus fermentum, Enterococcus faecalis, Staphylococcus aureus, Listeria monocytogenes*, and *Bacillus subtilis*], with genomic DNA from each species contributing to 12% of the community composition, and two yeasts (*Saccharomyces cerevisiae* and *Cryptococcus neoformans*) with genomic DNA from each contributing 2% to the community composition. The concentration of DNA was determined by a Qubit FlexTM fluorometer (Invitrogen, Carlsbad, CA) (**S1 Table**). In total, 80 samples were extracted using four kits.

### Library preparation and sequencing

The extracted DNA samples were prepared for Illumina next-generation sequencing using the NEBNext Ultra II FS DNA Prep kit from Illumina, part number E7805. The protocol was followed per the manufacturer’s instructions and the samples were fragmented for 15 minutes. Before library preparation, a fragmentation test was performed to determine the optimal fragmentation time. Post library QC was completed first by running the samples on the Agilent BioAnalyzer High Sensitivity Assay and then performing qPCR using the Kapa SYBR green assay for Illumina. Sequencing was conducted on the NovaSeq 6000, a full SP PE250 Illumina run type generating 9.9 × 10^8^ total reads.

### Bioinformatics

All sequence data were stored with unique de-identified subject IDs. The metagenomic profiles of each sample were characterized using both Kraken2 [8] version 2.1.1 and Metaphlan 3.0 [9] version 3.0.7 standard processing pipelines, both run with default parameter values; the standard Kraken2 database was used for Kraken2 classifications. These two pipelines were both applied to determine if, and how, analysis results might vary by algorithm; Kraken applies a *k*-mer based taxonomic classification approach, while Metaphlan bases its classifications on a catalog of clade-specific marker genes.

### Quality control

After initial bioinformatic processing, stacked bar plots of microbiome composition were assessed by site and kit (**Fig. 1**). Upon visual inspection, it was clear that one of the oral and vaginal swabs extracted using the HostZERO kit was incorrect: note, samples labeled “52V” and “52O” are shown in **Fig. 1** in their presumed correct positions; “52V” specifically appears to have an abundance of *H. parainfluenzae, R. mucilaginosa*, and *S. mitis*, all of which it shares in common with the other oral swab samples processed with the HostZERO kit. Taxa identified from the “52O” sample are primarily contaminants that were observed in multiple samples. It is likely these samples were swapped either at time of collection or during the extraction procedure. Therefore, during analysis, the samples were switched back to their presumed correct positions. We conducted our analysis with the sample excluded, in the likely swapped positions, and with the samples switched to their presumed correct position. Analysis results were consistent regardless of inclusion or exclusion of the samples. We chose to present the results with the samples in their presumed correct positions.

**Fig 1.**
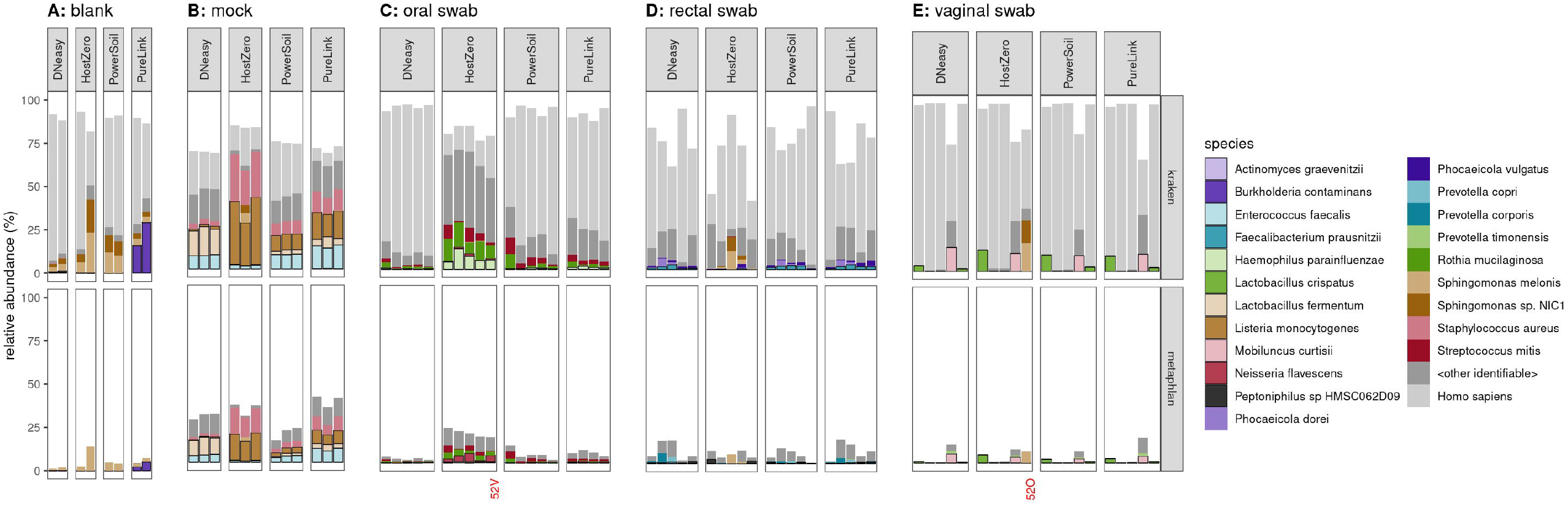
Fraction of reads assigned to *H. sapiens* versus microbial reads. Host reads are highly abundant among samples extracted with all kits. HostZERO does reduce the fraction of host reads (light grey), especially for the oral swabs better than other extraction kits. Some of the reads which are not mapped to any of the indicated taxa (including “other bacterial species”) remain unidentified by Metaphlan but may be identified as bacterial by kraken2; this is particularly true for the mock specimens.

### Statistical analysis

We measured alpha diversity using the Shannon and Chao1 indexes and observed species richness, using rarefied samples. Each alpha diversity index was compared between extraction kit types using the ANOVA test if the index was normally distributed, or the Kruskal-Wallis test if the index is non-normal. For comparing between test kits using paired within individual samples, beta diversity was measured using the Bray-Curtis distances and exploratory analysis with principal coordinate analysis (PCoA) plots or Non-metric Multi-dimensional Scaling (NMDS) was completed. To obtain a global overview of the sequencing results, nonmetric multidimensional scaling (NMDS) analysis was performed. We applied NMDS as implemented by the metaMDS function of the R package vegan [10] to the square-root-transformed and Wisconsin-standardized [11] matrix of species-level abundance estimates resulting from Metaphlan. To identify which extraction kit results best approximate the known composition of the mock communities, we used compositional bias analysis using the R package metacal [12]. Because the default bootstrapped metacal analysis method involves a log-transformation step, a small offset (or pseudocount) of 0.001 was added to the taxa abudance fractions before inputting them to metacal’s estimate_bias function; this leads to low but non-zero bias estimates when a given taxa is consistently undetected. We scored the ability of each kit to reproduce the pattern of relative abundances according to the geometric standard deviation—defined as the exponential of the standard deviation of log-transformed values, analogous to the geometric mean—of the bias values across all taxa. A metacal bias value of 1 for a given taxa indicates that there is no bias in relative abundance, while values above or below 1 indicate that the relative abundance of the taxa is over- or underestimated, respectively. The geometric mean of metacal bias values across all taxa is always unity since metacal aims to assess bias in relative, not absolute, abundance. A geometric standard deviation of 1 would indicate that the proportions of all taxa are in perfect accord with the input amounts, while higher values of the geometric standard deviation of bias values indicate bias values further away from the ideal value of 1. All statistical analyses were completed in R.

## Results and discussion

### Contamination

We identified known laboratory contaminants in our extraction blanks and mock community samples. Other groups have previously identified microbial communities that were associated with specific extraction kits or laboratory reagents often referred to as a “kitome” [13–16]. The most prevalent contaminant we identified was *Sphingomonas melonis*, which is a gram-negative bacterium from the genus *Sphingomonas* and is commonly found in soil, water, and plants [17]. *S. melonis* was detected in all extraction blanks, which contained sterile water. The contaminant was not present in all samples, although was identified in two mock communities and 11 clinical samples (four oral, six rectal, and one vaginal sample). *Burkholderia contaminans* were only detected in blanks and mock community samples that were extracted using the PureLink kit. *B. contaminans* were not detected in any of the clinical samples using any kit. Lastly, *Lactobacillus iners* and *Mobiluncus curtisii* were identified in two blank samples. The contaminants were in separate blank samples with very low abundance (0.07, 0.09 respectively). However, both samples containing these contaminants were extracted using the PureLink kit. Both bacteria are present in our clinical samples, and cross-contamination from clinical samples is possible.

### Mock community samples

The expected composition of the mock samples should be equal for eight bacterial species and two fungal species, with each representing 12% and 2% of genomic reads respectively. We also observed generally consistent results of microbial classification when using Kraken and Metaphlan to assign taxonomy. However, if investigators are interested in analyzing fungi, only the Metaplan database contains fungal sequences and should be used. All but one of the bacterial species—*Bacillus subtilis*— were observed in the data (**Fig. 2**). However, *Bacillus intestinalis* was observed in all mock samples and all kits, which were not supposed to be present in the mock community samples. Therefore, we suspected that *B. subtilis* was misclassified as *B. intestinalis* and included *B. subtilis* in the results in Figure 1. We then investigated the sequences included in the Metaphlan taxonomy database and discovered there were only two sequences covering 2.7 kb for *B. subtilis* and 86 sequences covering 79 kb included for *B. intestinalis* (by contrast, the Kraken database contains 307 sequences totaling 754 Mb for *B. subtilis* and only one sequence of 4 Mb for *B. intestinalis*). Sequences for the two species in the 16S region are 99% similar and may explain the misclassification, despite the WMS approach generally providing higher accuracy in microbial species classification [18,19]. This suggests that even when using WMS, there may be some species and strain level misclassification of microbial taxa that are closely related [20].

**Fig. 2.**
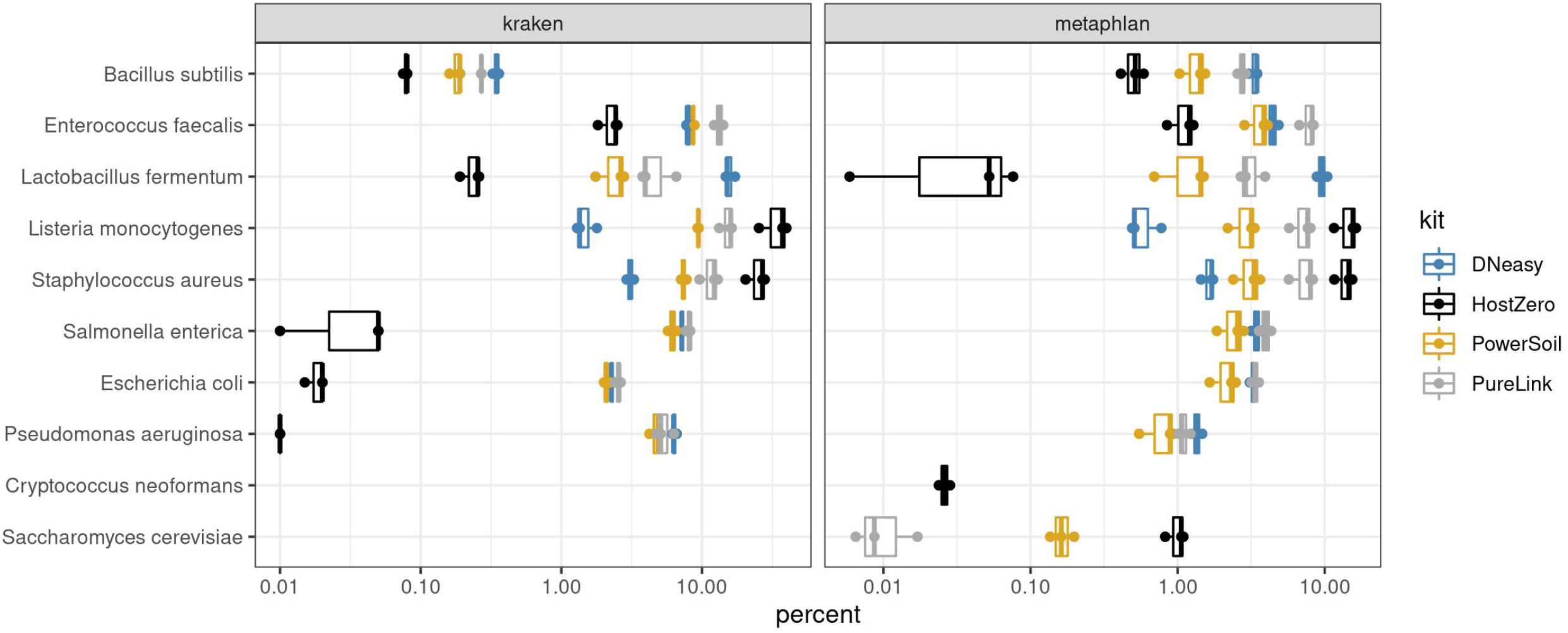
Percentage of mock community microbial reads extracted by kit. The composition of the mock samples from WMS analysis by the four DNA extraction kits. Eight bacterial species and two fungal species were expected to represent 12% and 2% of genomic reads respectively.

Consistent naming of taxa has become more complicated because the discovery of novel genera and species has grown exponentially due to increased use of metagenomic approaches in research. The International Code of Nomenclature of Prokaryotes delineates rules for the naming of Archaea and Bacteria. Validly published, continuously updated taxa names can be found on the List of Prokaryotic names with Standing in Nomenclature site (https://www.bacterio.net) [20,21]. The lack of integration of updated taxonomy into databases used in bioinformatics pipelines results in different groups reporting the same sequence as different species or levels of taxonomy, making it difficult to compare results across studies. Since most mock community samples with known percentages of bacteria or genomic DNA contain few species, investigators should periodically review changes in taxa nomenclature in addition to including analysis of mock communities with their clinical samples as a ground truth. When changes in the nomenclature of taxa commonly reported in the literature occur, it is important to note the previous name to allow for accurate comparison with previous studies.

Next, we determined that extraction kits differentially biased the percentage of reads attributed to microbial taxa. For example, *S. aureus* and *L. monocytogenes* were detected at the highest abundance using the HostZERO and PureLink kits, while *L. fermentum* was overrepresented when using the DNeasy method (**Fig. 3**). *S. cerevisiae* was detected at the highest level using the HostZERO kit. The other yeast species included in the mock community, *C. neoformans*, was only detected in samples extracted with the HostZERO kit. Both fungi included in the mock community were underrepresented in all mock community samples, with a median relative abundance for *C. neoformans* of 0.02% compared to 1.05% for *S. cerevisiae*, instead of the expected 2%. We also observed overrepresentation of *S. aureus* and *L. monocytogenes*, at the expense of representation of all gram-negative bacteria (*P. aeruginosa, E. coli, S. enterica)*. None of the gram-negative bacteria present in the mock communities were identified in the sequencing results of samples extracted using the HostZERO kit when analyzed using Metaphlan (**Fig. 2 & 3**). Interestingly, a small percentage of reads were assigned to the gram-negative bacteria when using Kraken to assign taxonomy for samples extracted with the HostZERO kit. Given that the gram negative bacteria that were present in higher abundance were correctly annotated when using Metaphlan and Kraken, it is likely because Metaphlan is very conservative relative to Kraken when assigning taxonomy to low abundance microbes [22]. Overall, Kraken has been shown to be a fast and accurate method. However, others have reported that features that allow for speed may result in more false positives when using Kraken, which can be remedied by filtering out low abundance reads prior to assigning taxonomy [23].

**Fig 3.**
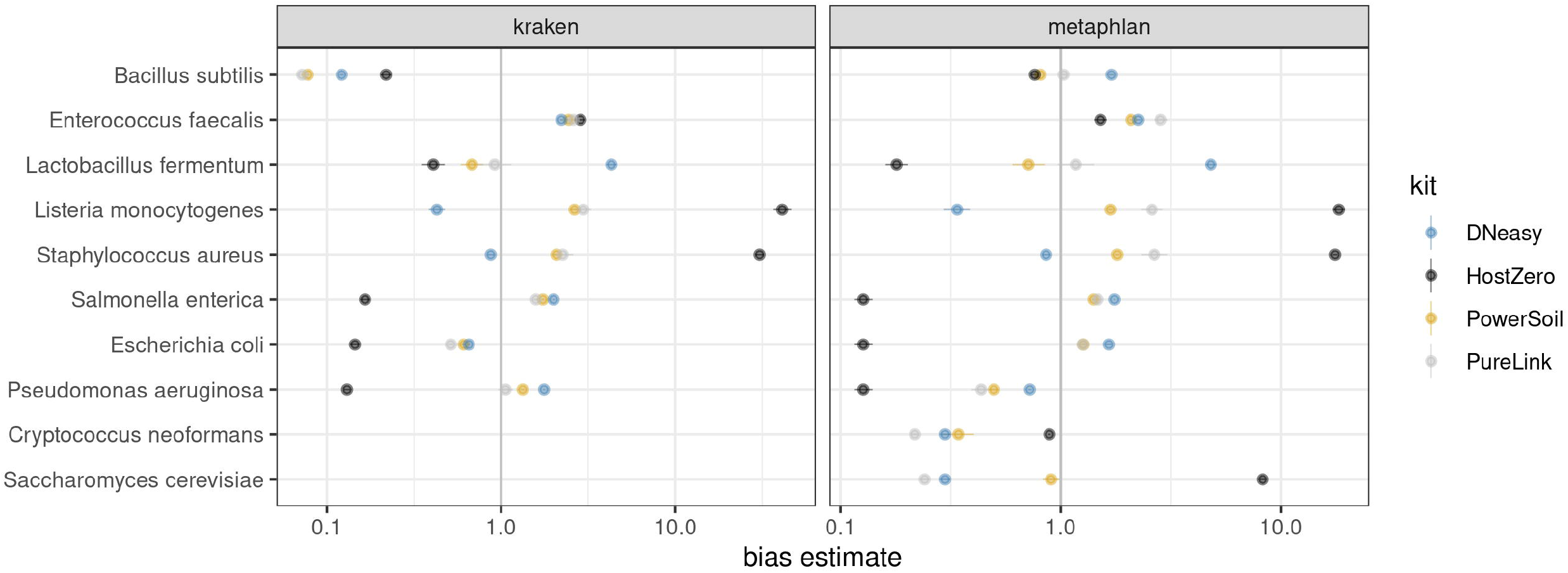
Metacal bias parameter estimates of mock communities. 1.0 indicates no relative bias for a particular species; values less than/greater than 1 indicate a bias toward lower/higher abundance values for the kit in question. The bias estimates are indicated on a log scale.

General differences between kit protocols are also likely contribute to the observed differences in mock community results. For example, HostZERO was designed to remove >90% of human reads from microbiome samples by degrading eukaryote DNA before the DNA purification step. Despite being designed to recover >85% of microbial DNA from samples, we found that the protocol was biased against gram-negative bacteria yet performed better for extracting DNA of fungi from mock communities than all other kits tested. However, compared to other kits, the relative abundance of bacteria within samples extracted using the HostZERO kit was distinctly different from the other kits.

The PureLink and PowerSoil Pro kits were both formulated to increase yields from difficult to lyse samples and had extraction results that were the most similar. Results for samples extracted using the PowerSoil Pro kit also best approximated the expected proportions of the mock communities, as quantified by compositional bias analysis using the R package metacal [12]; see **Table 2**. The geometric standard deviations of metacal bias value estimates suggest that extraction with the HostZERO kit yields the highest geometric standard deviations in both Kraken and Metaphlan-based analyses, owing to the considerably overestimated abundances of *S. aureus, E. faecalis*, and *S. cerevisiae* contrasting with the notably underestimated abundances of *S. enterica, E. coli, L. fermentum*, and *P. aeruginosa*.

**Table 2.** Dispersion of metacal bias estimates, as measured by geometric standard deviation (geo sd)

| Kit | Kraken geo sd | Metaphlan geo sd |
| --- | --- | --- |
| DNeasy | 3.09 | 2.60 |
| HostZERO | 11.2 | 7.62 |
| PowerSoil | 3.22 | 1.81 |
| PureLink | 3.37 | 2.60 |

### Clinical samples

The HostZERO kit yielded smaller fractions of reads assigned to *Homo sapiens* across all three sites sampled (oral, rectal, and vaginal) than did each of the other kits. When using HostZERO for extraction, in oral swabs, an average of 16.4% of reads in oral swabs were mapped to *H. sapiens* as compared to 85.4%, 73.6%, and 67.7% for DNeasy, PowerSoil, and PureLink, respectively; in rectal swabs, the HostZERO *H. sapien* mapping rate was 42.4% versus 60.5%, 66.9%, and 58.7% for the other kits; finally, in vaginal swabs, the rate for HostZERO was 74.5% versus 87.5%, 87.9%, and 83.3% (**Fig. 2**). The HostZERO-prepared samples had correspondingly higher fractions of reads assigned to bacterial taxa than did the samples prepared using other kits. For investigators interested in minimizing the number of human reads in their sequencing run, HostZERO could be the preferable kit. However, minimizing host reads also appears to bias the representation of microbial communities, as samples extracted using HostZERO look more distinctly different than differences between other extraction kits (**Fig. 4**). These differences are consistent with what we observed in the mock community samples.

**Fig. 4.**
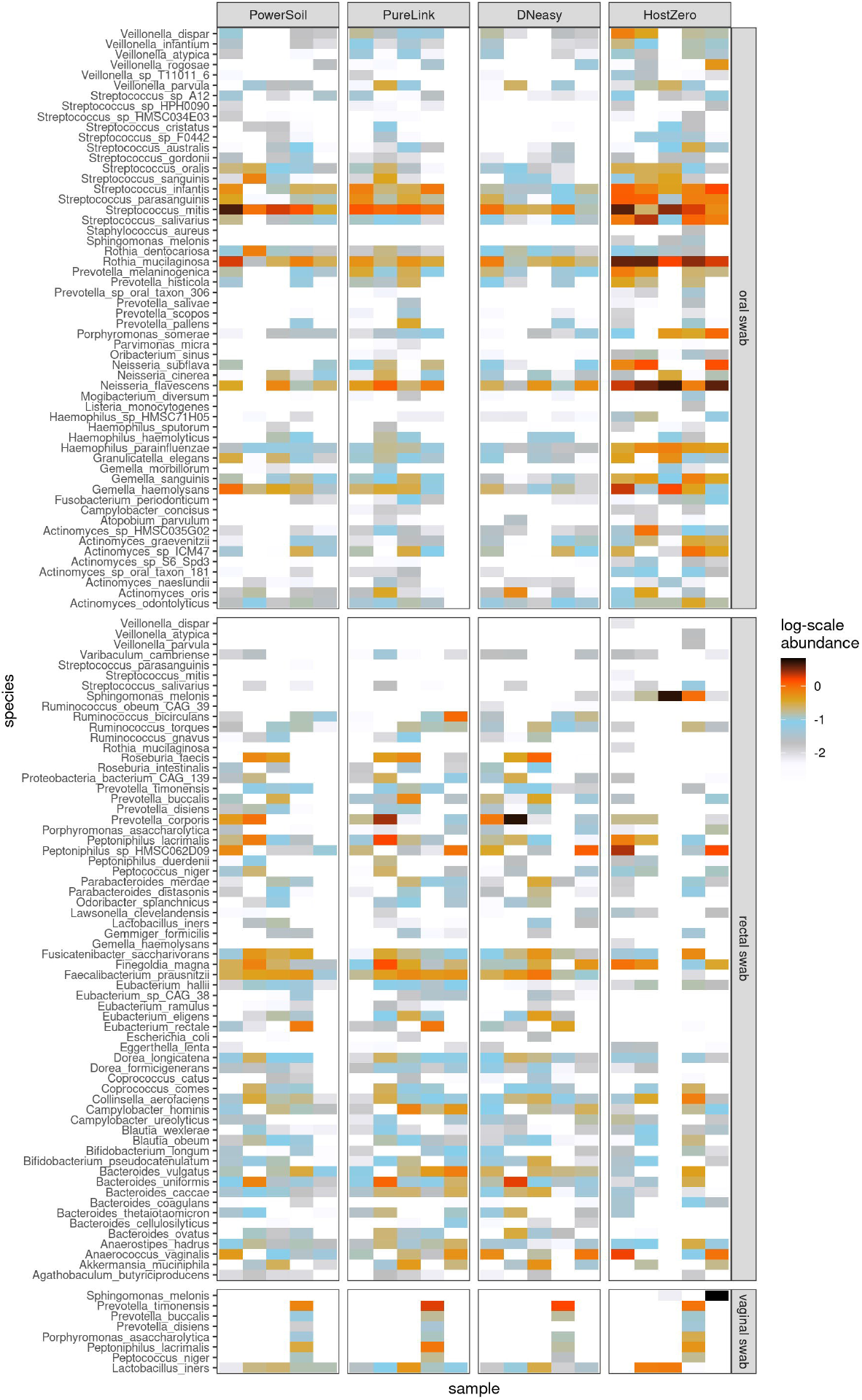
Top microbial species extracted for each clinical site by kit type. Five columns in each box represent the five samples from the volunteers.

However, as illustrated in **Figure 5**, when we assess the samples using NMDS the samples separate primarily by site, with the vaginal swabs exhibiting a more substantial dispersion than the other sites. Samples characterized as outliers (based on root-mean-square (RMS) displacement, a.k.a. Euclidean distance, from the corresponding within-kit-and-site NMDS coordinate centroid position exceeding a value of 2) are labeled in a “<kit>: <site>“ format on the NMDS plot of **Fig.5**: all four such samples were processed using the HostZERO kit, and three out of four were vaginal swab samples. Two of these outlier samples, including the lone non-vaginal (rectal) swab outlier, co-locate with the blank specimens.

**Fig. 5.**
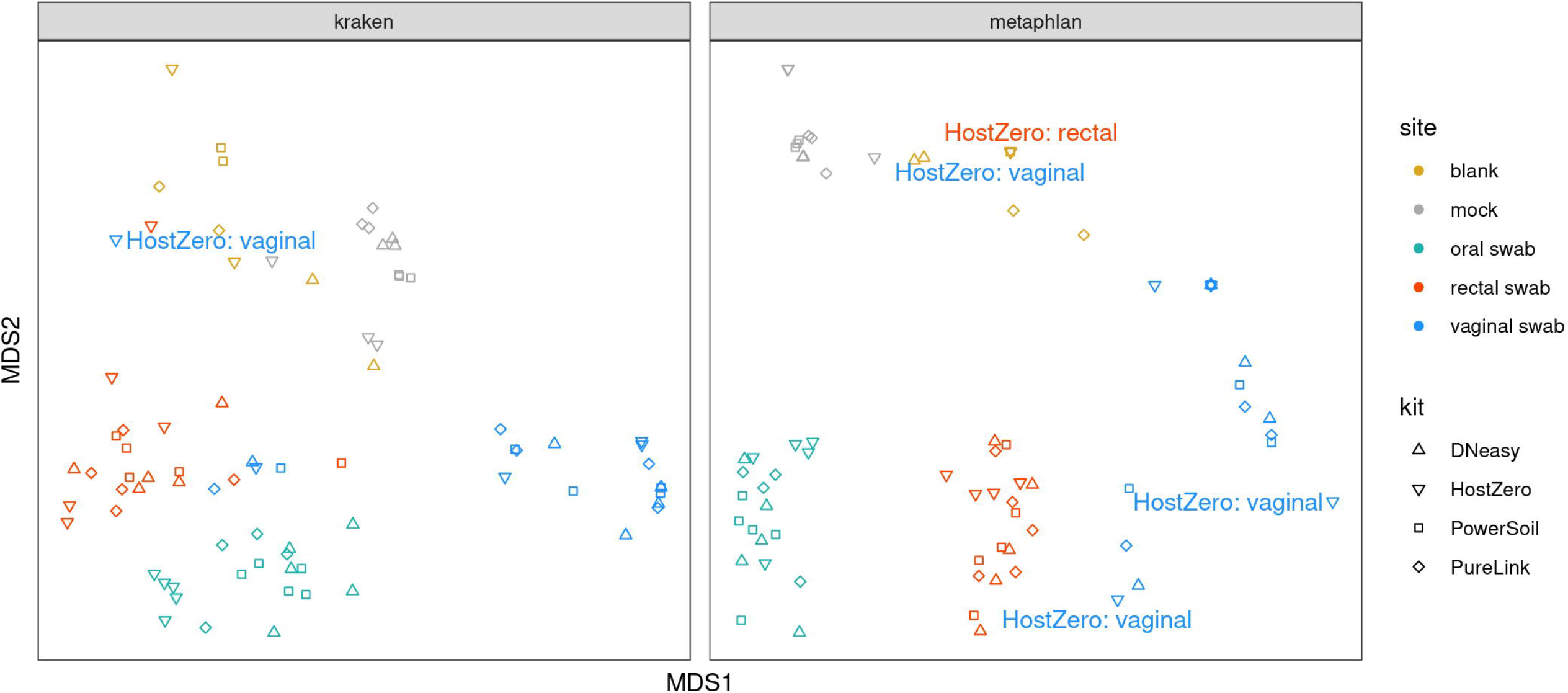
Sample Analysis Using Non-metric Multi-dimensional Scaling (NMDS). Samples cluster most tightly by type or site with HostZERO samples being most dispersed, particularly for vaginal samples.

The average RMS displacements of the samples from their corresponding group centroids within each kit/site combination are displayed in **Table 3**. All blank samples processed with the HostZERO and PowerSoil kits are assigned identical NMDS coordinates, leading to the 0 dispersion values shown in the first column of the table. For the mock samples, as well as both the rectal- and vaginal-site swabs, the HostZERO-processed sample sets have notably higher average RMS dispersions than any of the other kits. For the oral swabs, however, there is a much narrower range of dispersion values between kits, with the DNeasy-processed samples more dispersed than the HostZERO-processed samples. Overall, the communities represented by the HostZERO kit were most different from the other kits for rectal and vaginal sites and DNeasy community representation was most different for the oral microbiome samples. For investigators comparing microbiome communities across studies or conducting metaanalyses, the extraction method used may be an important covariate to include if there is variability across studies.

**Table 3.** Average root-mean-square (RMS) displacements from within-group centroids for two-dimensional NMDS coordinates by kit and site.

| Kit | Analysis | Blank | Mock | Oral swab | Rectal swab | Vaginal swab |
| --- | --- | --- | --- | --- | --- | --- |
| DNeasy | Kraken | 0.335 | 0.038 | 0.316 | 0.363 | 0.976 |
| HostZERO | Kraken | 0.65 | 0.382 | 0.113 | 0.729 | 1.429 |
| PowerSoil | Kraken | 0.044 | 0.035 | 0.174 | 0.532 | 0.875 |
| PureLink | Kraken | 0.233 | 0.062 | 0.306 | 0.352 | 1.01 |
| DNeasy | Metaphlan | 0.059 | 0.003 | 0.56 | 0.622 | 1.103 |
| HostZERO | Metaphlan | 0 | 0.613 | 0.492 | 1.203 | 1.907 |
| PowerSoil | Metaphlan | 0 | 0.031 | 0.401 | 0.526 | 0.913 |
| PureLink | Metaphlan | 0.439 | 0.176 | 0.465 | 0.46 | 1.039 |

Differences in the fractions assigned to fungal taxa or viruses appeared to be both smaller and less consistent across kits; this may be partially attributable to the lower overall abundance of these species, leading to reduced precision in the estimated abundance. For example, we did not detect any fungal taxa in our clinical samples. Although our mock communities did not include virus as a ground truth, we did identify some viral taxa in oral and rectal clinical samples for all extraction kits. Surprisingly, we also detected virus, predominantly *Staphylococcus* bacteriophage Andhra, in mock communities that did not contain virus using all kits except PowerSoil (**Fig. 6**). Additional studies to ground truth the mycobiome, virome and microbiome concurrently may be informative for investigators interested in studying the interaction across microbial communities.

**Figure 6.**
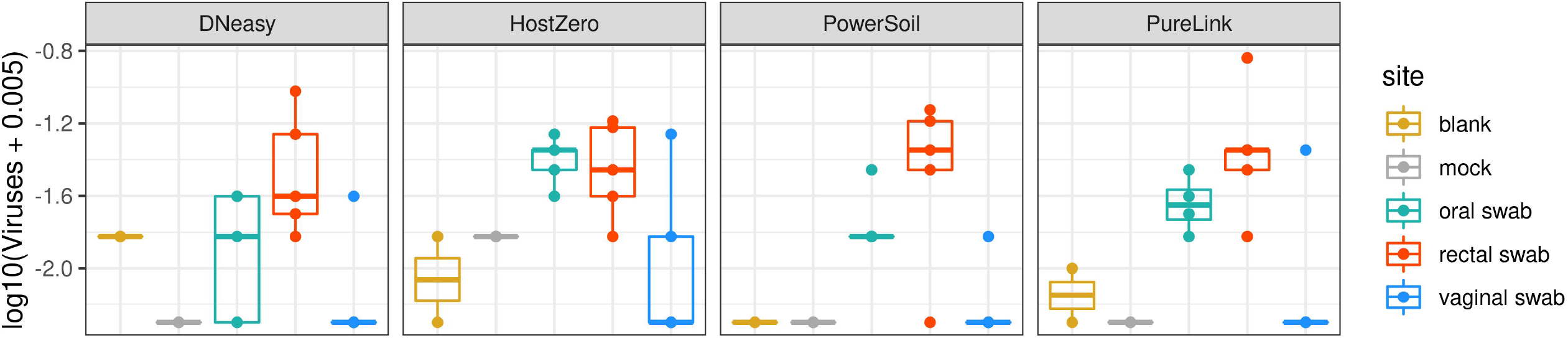
Total vial abundance for clinical samples by kit. Total viral (relative) abundance levels above are taken from Kraken2 analysis as viral sequences are not present in the Metaphlan database.

## Conclusion

Similar to reports of bias introduced by extraction kits when conducting amplicon-based sequencing, we observe bias in relative abundance of microbial taxa for WMS by extraction method. We observed very low levels of contamination that varied by kit and significant differences in relative abundance in mock community extractions by method. Although HostZERO was the only kit to extract both fungal species from the mock communities, it was at the expense of gram-negative bacteria. HostZERO was also superior in reducing the number of human reads present in a sample, although this may contribute to greater deviation from the expected yields when using this extraction kit. Given the biased introduced by extraction method, it is important for investigators conducting meta-analyses to include this as a covariate in their analyses when different methods are used across studies. Investigators should weight the benefits and biases introduced by different methods to minimize influences to their analysis.

## Acknowledgments

DNA library prep and sequencing were performed by the Genomic Sequencing and Analysis Facility at UT Austin, Center for Biomedical Research Support. RRID#: SCR_021713.

## Supporting information

### S1 Table

Concentrations of extracted DNA.

### Data and code availability

Sequence data in the process of being uploaded to SRA and code is available at https://github.com/mlwright97/WMSextraction.

